# TGFβ mediated structural remodeling facilitates optic fissure fusion and the necessity of BMP antagonism in this process

**DOI:** 10.1101/131599

**Authors:** Max D. Knickmeyer, Juan L. Mateo, Priska Eckert, Eleni Roussa, Belal Rahhal, Aimee Zuniga, Kerstin Krieglstein, Joachim Wittbrodt, Stephan Heermann

**Author notes:** present address: School of Medicine and health sciences, An-Najah National University, Nablus, Palestine (Belal Rahhal). Author for correspondence: Stephan Heermann.

## Abstract

The optic fissure is a transient gap in the developing vertebrate eye, which must be closed as development proceeds. A persisting optic fissure, coloboma, is a major cause for blindness in children. Multiple factors are genetically linked to coloboma formation. However, especially the process of optic fissure fusion is still largely elusive on a cellular and molecular basis.

We found a coloboma phenotype in mice with a targeted inactivation of the transforming growth factor 2 (TGFβ2). Here the optic fissure margins got in touch, however, failed to fuse. Transcriptomic analyses indicated TGFβ mediated ECM remodeling during optic fissure fusion. For functional analyses, we switched model systems and made use of zebrafish. We found TGFβ ligands expressed in the developing zebrafish eye, and the ligand binding receptor in the optic fissure. Using a new in vivo TGFβ signaling reporter, we also found active TGFβ signaling in the margins of the optic fissure. We addressed the function of Cadherin 6 (*cdh6*), one of the TGFβ regulated genes, by knock down experiments in zebrafish and found a prominent coloboma phenotype. *Cdh6* was often found involved in processes of epithelial to mesenchymal transition (EMT), strengthening our hypothesis that an EMT-like process is also necessary for optic fissure fusion. Furthermore, we found Gremlin 2b (*grem2b*) and Follistatin a (*fsta*), homologs of TGFβ regulated bone morphogenetic protein (BMP) antagonists, expressed in the optic fissure margins, indicating the necessity of a localized inhibition of BMP signaling. Finally, we show that induced BMP expression is sufficient to inhibit optic fissure fusion. Together with our previous findings this indicates a dual role of BMP signaling during optic fissure closure.

**Summary statement:** TGFβ is crucial for optic fissure fusion, involving *cdh6*. TGFβ mediated optic fissure fusion is potentially hampered by BMP signaling, which is blocked by TGFβ induced BMP antagonists within the optic fissure margins.

## Introduction

The optic fissure is a transient gap in the ventral domain of the developing vertebrate eye. During a specific period of development, it is an entry route used by cells of the periocular mesenchyme and embryonic vasculature to enter the eye. However, it is necessary that the fissure is closed as development proceeds. A persisting fissure is termed coloboma. A coloboma can affect vision severely and is a frequent cause for blindness in children (Onwochei et al., 2000). Many genes and signaling pathways have been linked to coloboma formation (Graw, 2003, Westenskow et al., 2009, Bankhead et al., 2015, Chen et al., 2013, Cai et al., 2013, Miesfeld et al., 2015, Matt et al., 2008, Lupo et al., 2011, Lee et al., 2008, Heermann et al., 2015), resulting in a growing gene coloboma network (Gregory-Evans et al., 2004, 2013). Coloboma often is part of a multi organ syndrome, e.g. CHARGE syndrome or renal coloboma syndrome, linked to Pax2 and Chd7 respectively (Torres et al., 1996, Favor et al., 1996, Sanlaville and Verlores, 2007, Fletcher et al., 2006, Bower et al., 2011). Notably, the morphology of coloboma phenotypes is highly variable. Defects or alterations in some of these signaling pathways, for example, result in vast extended coloboma (Bankhead et al., 2015, Miesfeld et a., 2015), likely originating from morphogenetic defects. Such an early morphogenetic defect, resulting in coloboma, was demonstrated recently by a precocious arrest of the neuroretinal flow during optic cup formation, induced by ectopic expression of a BMP ligand (Heermann et al., 2015). Importantly, such massive coloboma phenotypes are different from colobomata resulting from a hampered fusion process of the optic fissure margins. Within these margins, the prospective neuroretina and the prospective retinal pigmented epithelium (RPE) share a basement membrane. With the onset of fissure fusion, the prospective neuroretina and RPE are separated and after that, reside on their own basement membrane respectively. This fusion of the two opposing fissure margins and the separation of the neuroretina from the RPE are demanding major structural remodeling. This includes the dissolution of the basement membrane within the margins (James et al., 2016), the loosening of cell-cell contacts in between prospective neuroretinal cells and RPE precursors and eventually establishment of new connections within the prospective neuroretina and RPE, respectively. N-cadherin and α-catenin were shown to be important in this context (Masai et al., 2003, Chen et al., 2012). Although the overall process that has to be executed is known quite well, only very little is known about how the actual fusion of the opposing optic fissure margins is achieved on a cellular and molecular basis. TGFβ is a well-known modifier of the extracellular matrix (ECM), in various processes during development and disease (Wu and Hill, 2016, Zhang et al., 2016, Thiery et al., 2009, Mercado-Pimentel and Runyan, 2007, Johansson et al., 2013). Here we address the role of TGFβ signaling for optic fissure fusion, making use of mouse (*Mus musculus*) and zebrafish (*Danio rerio*).

## Results and Discussion

### Loss of TGFβ2 results in coloboma

In mouse, three TGFβ isoforms are encoded in the genome (TGFβ1, 2 and 3). Targeted inactivation of TGFβ2 results in several phenotypes, also affecting the eye (Sanford et al., 1997), e.g. a remaining primary vitreous, a Peters anomaly like phenotype and an altered neuroretinal layering. In addition to these phenotypes, we identified a persistent optic fissure in TGFβ2 mutant embryos (Figure 1B, see A as control). TGBβ2 dependent coloboma were first discovered in TGFβ2/GDNF double mutants, (Rahhal, Heermann et al., 2009 unpublished observations) and subsequently also in TGFβ2 single mutants (this study) but not in GDNF single mutants. Notably, no phenotype affecting eye development was described in any of the three independently generated GDNF mutant mouse lines (Pichel et al., 1996, Moore et al., 1996, Sanchez et al., 1996), though GDNF expression in the developing eye was documented (Hellmich et al., 1996). Therefore, the coloboma phenotype must be assigned to TGFβ2 function. Notably, in the TGFβ2 mutants the optic fissure margins were able to get in close proximity to each other, but ultimately failed to fuse and instead grew inwards (Figure 1B). TGFβ signaling is also necessary for the fusion of the palatal shelves during development (Sanford et al. 1997, Proetzel et al., 1995). There, the ligands TGFβ2 and TGFβ3 have slightly different functions. While in mice with targeted inactivation of TGFβ2, the palatal shelves stay apart and a gap remains (Sanford et al., 1997), in TGFβ3 mutant mice the palatal shelves touch, but do not fuse (Proetzel et al., 1995, Taya et al., 1999). The latter scenario is reminiscent of what we observed for optic fissure closure in TGFβ2 mutant mouse embryos. The fissure margins meet, but do not fuse (Fig. 1B, see A as control).

**Figure 1:**
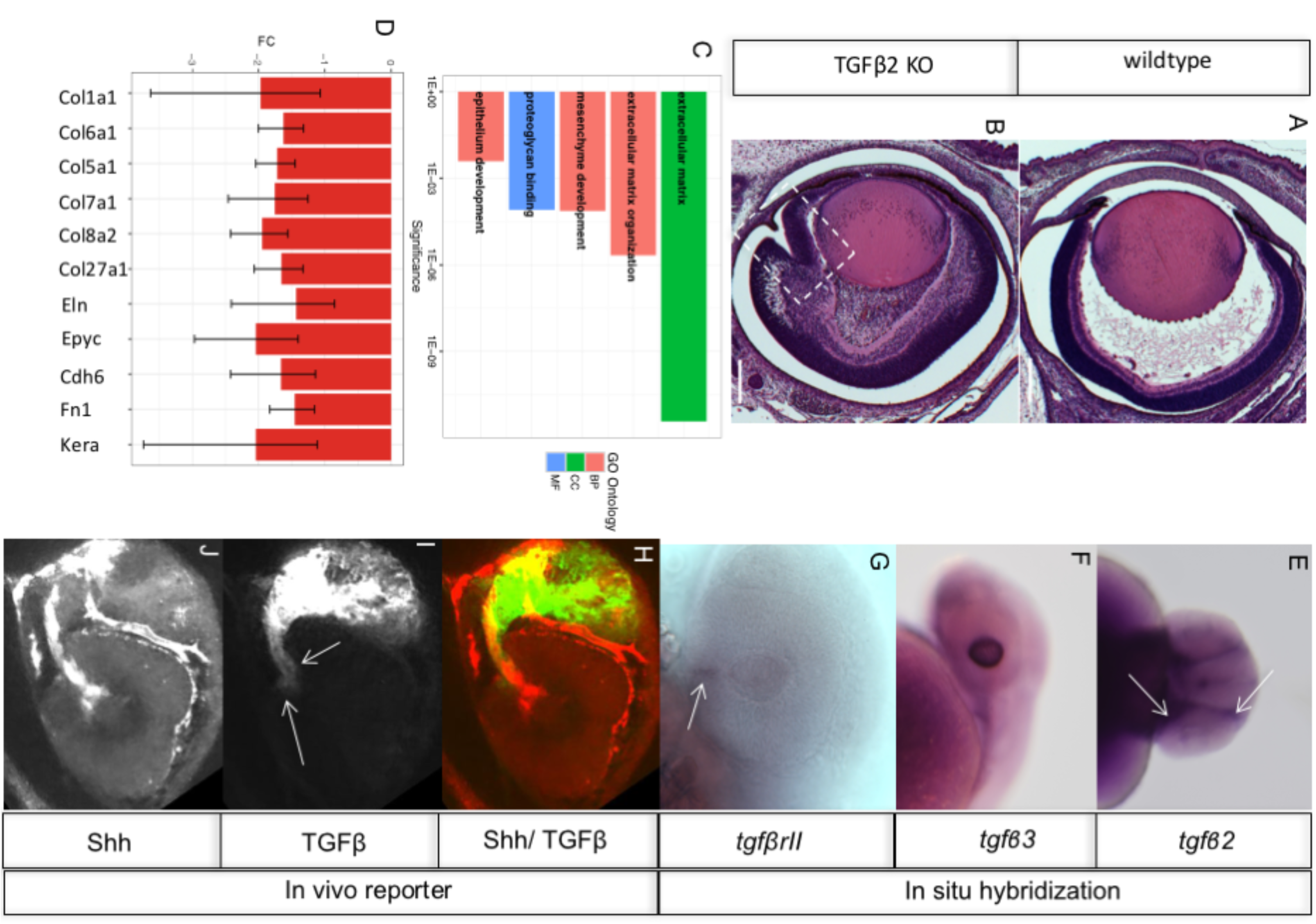
Loss of TGFβ2 ligand results in coloboma, ECM remodeling is affected and TGFβ ligands and TGFβ signaling in the developing zebrafish eye (**A**) Frontal section of a wildtype embryo (E18.5) as control (H&E staining). (**B**) Frontal section of a TGFβ2 KO embryo (E18.5) (H&E staining). Note the persisting optic fissure (boxed). Importantly the fissure margins must have met, but grew inwards, rather than fusing. Scale bars represent 200μm. (**C**) Selected terms enriched in the set of down-regulated genes in TGFb2/GDNF KO microarray based on the analysis with the tool gProfiler. BP=Biological Process, CC=Cellular Component, MF=Molecular Function. SP_PIR_KEYWORDS=Swiss-Prot Protein Information Resource. (**D**) Level of misregulation for several ECM related genes represented as fold change (FC). Error bars represent the 95% confidence interval. (**E**) in-situ hybridization of *tgfb2* (30 hpf), frontal view. Note expression of *tgfb2* in the periocular mesenchyme (arrows). (**F**) in-situ hybridization of *tgfb3* (30 hpf), lateral view. Note expression of *tgfb3* in the distal part of the developing lens. (**G**) in-situ hybridization of *tgfβrII* (30 hpf), lateral view. Note expression of *tgfβrII* in the optic fissure region (arrow). (**H**) combination of *in vivo* signaling for TGFβ (green) and Shh (red) split into the single reporter for TGFβ (**I**) and Shh (**J**). Note the active TGFβ signaling in the optic fissure margins (**I**, arrows).

### Ocular expression of ECM genes during optic fissure fusion is TGFβ dependent

Many genes have been linked to coloboma formation (Gregory-Evans et al., 2004, 2013), however, the structural changes occurring at the optic fissure margins during fusion are still not well understood. At the time when the margins of the optic fissure get in touch, structural changes inside these margins are essential to occur in order to allow their fusion. The basal membrane must be dissolved (James et al., 2016) and the epithelial structure must be loosened and re-established after fusion, involving the setup of new cell-cell and cell-matrix connections. We propose that this process occurs in an “epithelial to mesenchymal transition (EMT) like” process. Here, after an initial disassembly and consecutive fusion of the fissure margins, two new epithelia are established. These new epithelia, the neuroretina and the RPE, each contain a separate basal membrane. For this process to occur, the ECM has to be remodeled intensively. TGFβ signaling is well known for its ECM remodeling activity in various processes (Wu and Hill, 2016, Zhang et al., 2016, Thiery et al., 2009, Mercado-Pimentel and Runyan, 2007, Johansson et al., 2013). We thus addressed the potential transcriptional ECM regulation during optic fissure closure using our coloboma model. We quantified the levels of mRNAs from E13.5 embryonic eyes using Agilent microarray chips. To this end we compared RNA, harvested from eyes of wildtype as well as TGFβ2/GDNF double mutant embryos, from which the coloboma phenotype was assigned to TGFβ2 function only (see above). Henceforth the effects are termed TGFβ dependent. We processed the obtained microarray data, focusing on the genes significantly down-regulated. Performing a functional analysis of these genes we found as most prominent terms ECM, ECM organization, mesenchyme development, epithelium development or proteoglycan binding (Figure 1C). Next, we analyzed the expression levels of distinct ECM genes (Figure 1D). We found reduced expression of collagen genes Col1a1, Col6a1, Col5a1, Col7a1, Col8a2 and Col27a1 in combination with reduced levels of Elastin (Eln) and Epiphycan (Epyc), pointing towards reduced levels of fibrillogenesis in the extracellular space. Reduced levels of Cadherin 6, a factor often linked to EMT (Gugnoni et al., 2017, Clay and Halloran, 2014), strengthened our hypothesis of an EMT-like process and reduced levels of Fibronectin 1, indicated affected cell-matrix contacts. Furthermore, we found reduced levels of Keratocan (Kera), which was shown to be involved during TGFβ2 regulated corneal development (Saika et al., 2001).

### TGFβ ligands, the ligand binding receptor and TGFβ signaling in zebrafish optic cups

Next, we wanted to functionally address identified target genes with respect to their necessity for optic fissure fusion. The zebrafish model is well-suited to address genes in a loss of function context. In addition, zebrafish embryos are easily accessible for analysis, due to their extrauterine development and partial transparency. To assure that a switch of model system to zebrafish is feasible, we first investigated the expression of TGFβ ligands and the TGFβ ligand binding receptor during zebrafish eye development. We found *tgfβ2* expressed mainly in periocular tissue (Figure 1E) whereas *tgfβ3* was expressed mainly in the developing lens (Figure 1F). The ligand binding receptor *tgfβrII* we found expressed at the site of the optic fissure (Figure 1G). The expression of TGFβ ligands in the eye and especially the TGFβ ligand binding receptor within the optic fissure margins suggests that TGFβ is also involved in zebrafish optic fissure fusion. We next became interested in the activity of TGFβ signaling. To assess the dynamics of activated TGFβ signaling *in vivo* during zebrafish development, we established a transcriptional TGFβ sensor in a stable transgenic line. The reporter system is based on Smads, the canonical transcription factors transducing TGFβ signaling (Heldin et al., 1997). We used repetitive Smad Binding Elements (SBE) from the human plasminogen activator inhibitor (PAI). Such a reporter has been intensively used for years as a luciferase assay to assess the amount and activity of TGFβ in cell culture (Dennler et al., 1998) and in mice (Lin et al., 2005). The repetitive SBEs were combined with a minimal promoter element to drive the expression of a membrane bound GFP (GFPcaax) (Figure S1A). We then established a stable transgenic zebrafish line. Activated TGFβ signaling can be observed during development, e.g. by domains in the forebrain region as well as the distal tail (Figure S1B). We validated the functionality of this reporter line employing the established TGFβ signaling inhibitor SB431542 (Inman et al., 2002, Laping et al., 2002). We observed a drastic reduction in reporter activity after treatment with SB431542 (Figure S1C). Next, we wanted to know whether TGFβ signaling was active in the optic fissure margins. We employed the TGFβ signaling reporter line in combination with a reporter for Shh signaling (analog to Schwend et al., 2010) to relate the fissure to the optic stalk and performed *in vivo* time-lapse analyses. We found the TGFβ signaling reporter active within the optic fissure margins (Figure 1H and I see J for shh reporter activity only). Taken together our data indicate that TGFβ signaling is indeed active in the optic fissure margins of the zebrafish.

### Cadherin 6 is essential for optic fissure fusion

Next, we aimed at addressing the functional implications of the TGFβ regulated ECM genes during optic fissure fusion in zebrafish. Among the set of identified TGFβ regulated target genes, we found Cdh6. Cdh6 was shown to be involved in processes of epithelial to mesenchymal transition (EMT) in development and disease (Gugnoni et al., 2017, Clay and Halloran, 2014, Sancisi et al., 2013) and was identified to be TGFβ dependent in thyroid cancer (Sancisi et al., 2013). During EMT, epithelialized cells exit the united cell structure and obtain mesenchymal features. We hypothesize that a part of the optic fissure fusion program is EMT-like, in which the basement membrane is degraded (James et al., 2016) and the cells of the optic fissure margins establish the contact between the margins (Eckert et al., in preparation). However, the process is somewhat reversed consecutively, because soon after the fusion is established, two separated epithelia, residing on an own basement membrane respectively, are generated, the neuroretina and the RPE. We found *cdh6* expressed within the optic fissure margins (Figure 2A-C). We subsequently made use of a Morpholino oligonucleotide (MO) induced knock down approach using established MO sequences (Liu et al., 2008, Kubota et al., 2007).

**Figure 2:**
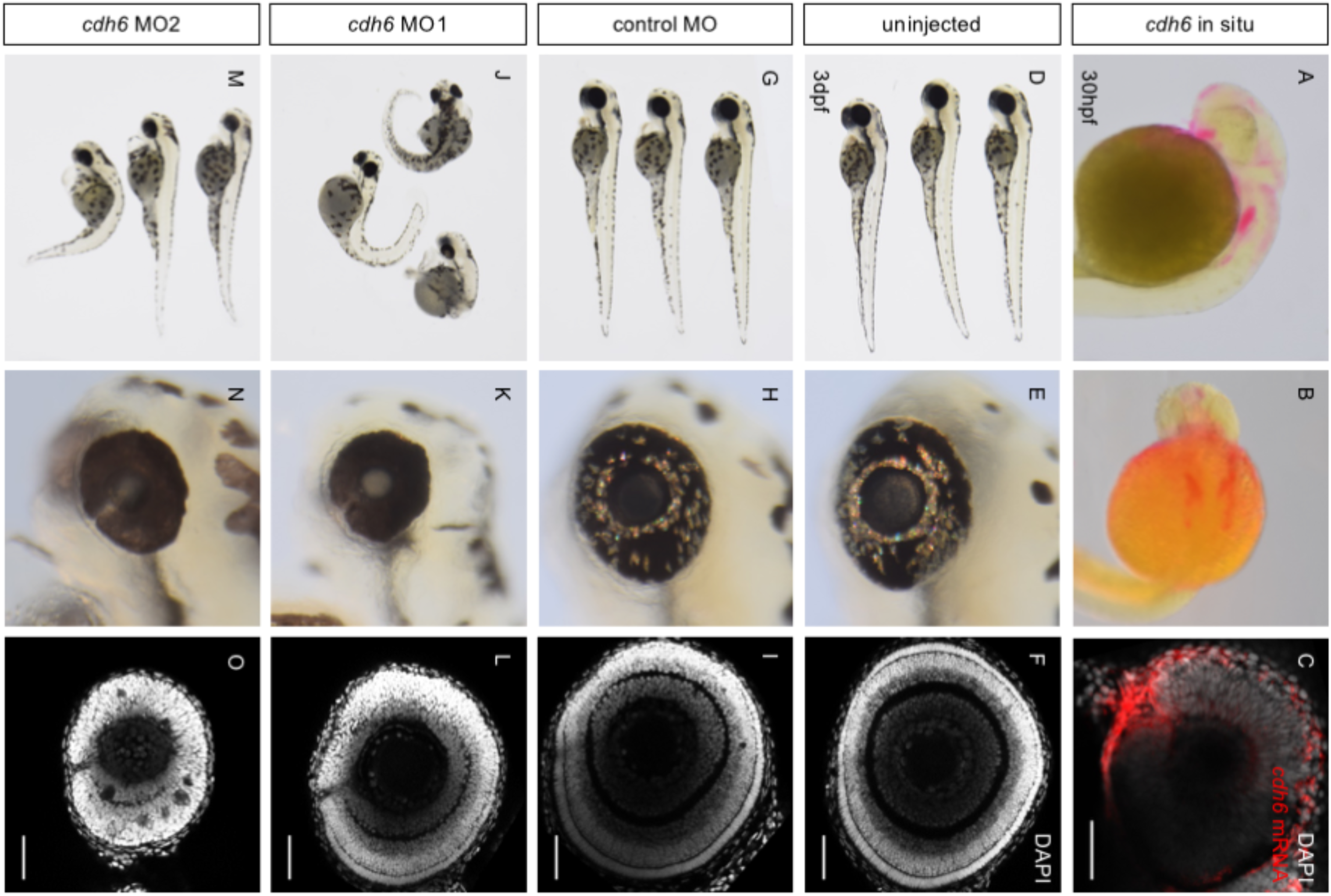
Knockdown of *cdh6* causes coloboma in zebrafish. (A, B) Whole mount in situ hybridization for *cdh6* in lateral (A) and dorsal perspective (B), please note the *cdh6* expression within the optic fissure (C) confocal image of *cdh6* whole mount in situ hybridization, (D, G, J, M) gross morphology of control embryos and morphants at 3dpf. (E, H, K, N) lateral view of the eyes of control embryos and morphants at 3dpf. *cdh6* morphants show coloboma, decreased eye size. (F, I, L, O) whole-mount eyes of control embryos and morphants at 3dpf, stained with DAPI. In addition to coloboma, *cdh6* morphants also show disrupted retinal layering (O). scale bars represent 50μm, orientation: nasal to the left

By screening with a stereomicroscope, we found that MO induced knockdown of *cdh6* caused coloboma in 31.6% (11.5ng of MO1) to 50.6% (4.5ng of MO2) of injected embryos (Figure 2J-L, M-O, Table, see 3D-I as control). Additional 20-32.9% of morpholino injected embryos showed an abnormally pigmented ventral retina, possibly indicating coloboma. 25.3% of *cdh6* MO1 injected animals showed severe defects of gross morphology and the eyes. Embryos injected with a control MO as well as uninjected animals only sporadically showed abnormal pigmentation or severe defects. Optical sections performed by confocal imaging of embryonic eyes with a nuclear staining confirmed the persistence of the optic fissure in *cdh6* morphants (Figure 2L, O). We also observed, in accordance with previous findings (Liu et al., 2008), that *cdh6* morphant eyes are reduced in size and show delayed and defective retinal differentiation.

### TGFβ mediated BMP antagonism during optic fissure closure

It is well-established that BMP4 in cooperation with Vax2 is important for the definition of cellular identities along the dorsal ventral axis within the vertebrate eye (Koshiba-Takeuchi et al., 2000, Sasagawa et al., 2002, Mui et al., 2005, Behesti et al., 2006). In line with these data we found *bmp4* expressed dorsally and *vax2* expressed ventrally within the zebrafish optic cup (Figure 3A and B). Notably, BMP signaling is also important for other aspects of eye development, like the optic fissure generation (Morcillo et al., 2006) and the optic cup formation itself (Heermann et al., 2015). In the latter study, it was shown that an inhibition of BMP signaling is crucial for a bilateral neuroepithelial flow to occur over the distal rim of the developing optic cup. This BMP signaling inhibition is achieved by the BMP antagonist Follistatin a (*fsta*). We found two antagonists for BMP signaling, Follistatin and Gremlin 1, transcriptionally down-regulated in our murine coloboma model (Figure 3C). Furthermore, we found homologous genes (*grem2b* and *fsta*) expressed in the optic fissure margins of zebrafish (Figure 3D and E). BMP4 is a secreted ligand and can potentially diffuse and act over extended distances. The expression of the two BMP antagonists in the optic fissure hints at a functional requirement to locally suppress BMP activity in this domain. Above we showed that TGFβ is relevant for ECM remodeling during optic fissure fusion and overall TGFβ signaling is well-known for its ECM remodeling activity in various processes (Wu and Hill, 2016, Zhang et al., 2016, Thiery et al., 2009, Mercado-Pimentel and Runyan, 2007, Johansson et al., 2013). Notably, BMP signaling seems to counteract these TGBβ induced changes often (Izumi et al., 2006, Zeisberg et al., 2003, Wang and Hirschberg, 2003, 2004). We propose that TGFβ induced local BMP antagonism is unleashing the ECM remodeling capacity of TGFβ (Figure 4F, scheme). This concept is new for optic fissure fusion, however, was shown to be accurate in the context of glaucoma (Zode et al., 2009). The BMP antagonist Gremlin has been linked to cleft lips in humans (Al Chawa et al., 2014) indicating that the level of BMP signaling is also tightly controlled during fusion processes there. We suggest that the combined expression of two BMP antagonists is important to provide robustness to the system. BMP antagonists are often expressed redundantly (Khokha et al., 2005, Eggen and Hemmati-Brivanlou 2001, Stottmann et al., 2001) pointing at the importance of their function. According to this it is not surprising that the loss of a single BMP antagonist does not result in coloboma (Grem1 knock out, not shown).

**Figure 3:**
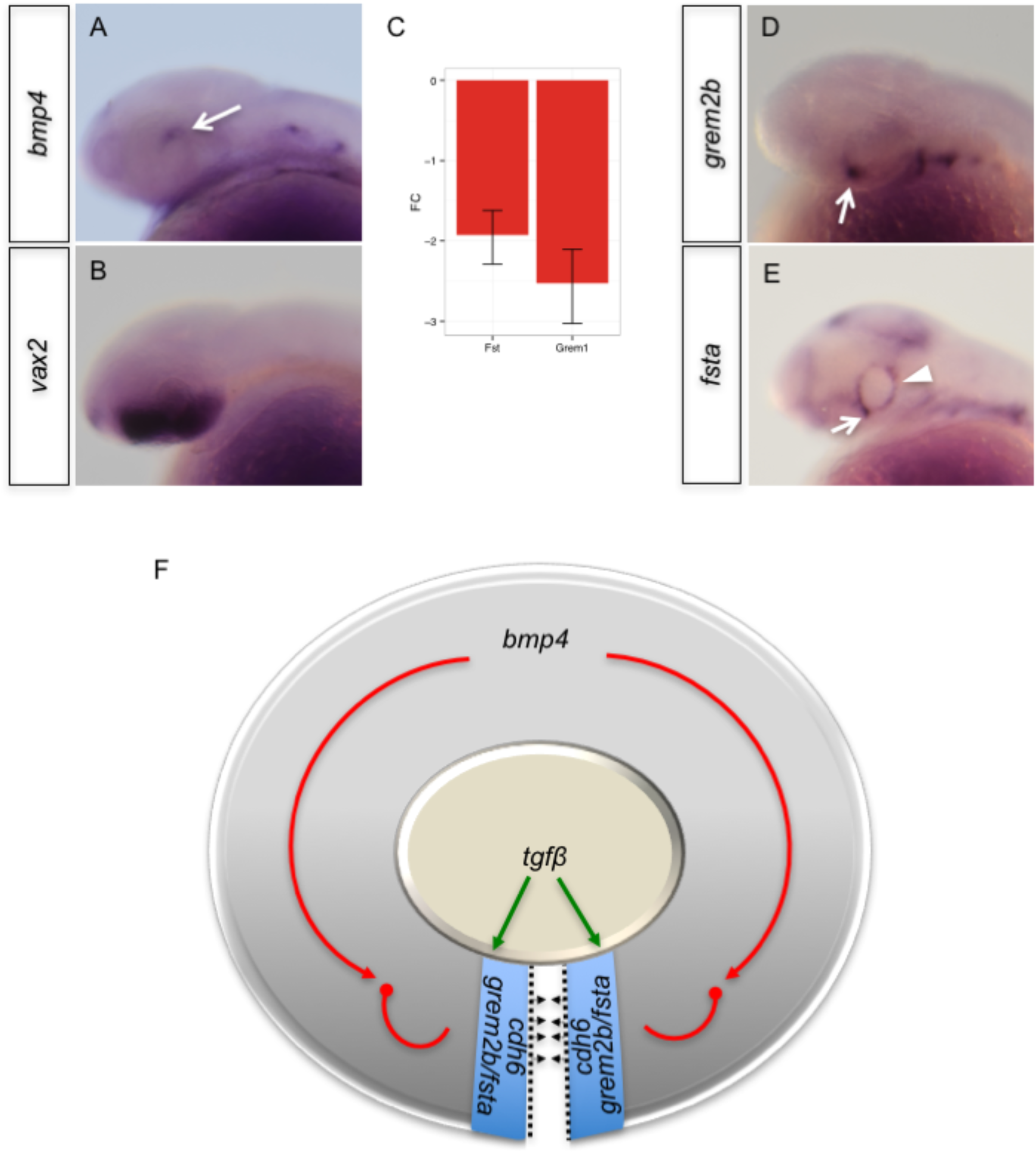
BMP antagonists are expressed in the optic fissure margins opposing the dorsally expressed BMP4, TGFβ interplay in optic fissure closure (scheme) (**A**) in-situ hybridization of BMP4 (30 hpf), lateral view. Note the localized dorsal expression of *bmp4* (arrow) within the optic cup. (**B**) in-situ hybridization of *vax2* (30 hpf), lateral view. Note the localized ventral expression of *vax2* within the optic cup. (**C**) Expression analysis of gremlin and follistatin, both being BMP antagonists, revealed a decrease in TGFb2 KO (TGFb2/GDNF KO) as represented by the fold change (FC). Error bars represent the 95% confidence interval. Corrected p-value of control gene expression compared to KO for follistatin and gremlin, 1.2E-3 and 5.5E-3 respectively. (**D**) in-situ hybridization of *grem2b* (30 hpf), lateral view. Note expression of *grem2b* in the optic fissure margins (arrow). (**E**) in-situ hybridizations of *fsta* (30 hpf), lateral view. Note expression of *fsta* in the optic fissure margins (arrow) and the developing CMZ (arrowhead). **(F)** TGFβ is inducing the expression of BMP antagonists, expressed in the optic fissure margins. We propose that this local inhibition of BMP signaling unleashes the capacity of TGFβ to remodel the ECM, enabling optic fissure fusion.

We next wanted to functionally test our hypothesis, that localized BMP inhibition in the optic fissure margins is necessary for TGFβ induced ECM remodeling and consecutive optic fissure fusion. To this end, we generated a transgenic line allowing for heat shock inducible *bmp4* expression (*tg(hsp:bmp4 cmlc2:GFP)*). With the help of this transgenic line we aimed at an oversaturation of the BMP antagonists and test whether or not this affects optic fissure fusion. Since the morphogenesis of the optic cup is dependent on BMP antagonism (Heermann et al., 2015), the timing of the heat shock induced expression of *bmp4* was critical. Thus, we first tested the outcome of heat shock induced *bmp4* expression at different successive stages of development (Figure S2, Figure 4). Induced expression at 21hpf and 22hpf resulted in a vast coloboma phenotype (Figure S2 A-A’’ and B-B’’, see D-D’’ and E-E’’ as control), well in line with the coloboma observed in our previous analyses (Heermann et al., 2015). This indicates that the transgenic line is functional but it also indicates that the onset of induction was too early and was affecting optic cup morphogenesis. The induced expression of *bmp4* at 23hpf resulted in a milder coloboma phenotype, with less affected cup morphogenesis (Figure S2 C-C’’, see F-F’’ as control). Notably, the resulting coloboma phenotype from induced *bmp4* expression at 24hpf, 25hpf and 26hpf was comparable, not showing defects in optic cup morphogenesis (Figure 4 A-C’’, see D-H’’ as control). According to our hypothesis (Figure 3F, scheme), active BMP signaling would suppress TGFβ induced structural remodeling within the optic fissure margins. In order to test this hypothesis further, we investigated *cdh6* expression in control embryos and heat shock induced *bmp4* expressing embryos. Indeed, the *bmp4* expression resulted in reduced levels of *cdh6* within the zebrafish eye (Figure 4 I, J, see K, L as control), indicating that our hypothesis is accurate.

**Figure 4:**
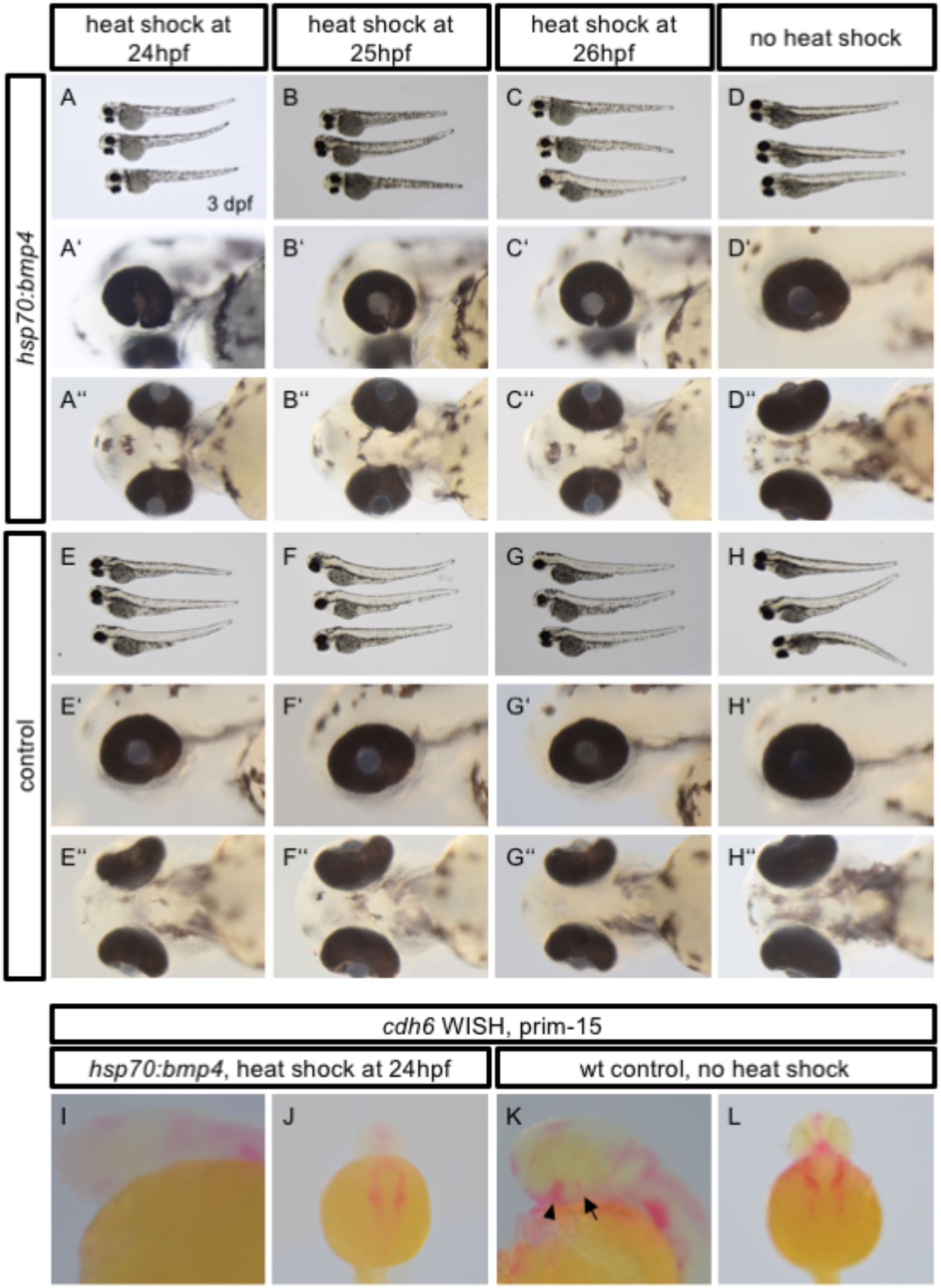
*bmp4* inhibits optic fissure closure and decreases expression of - regulated *cdh6*. (**A-H**) Gross morphology of (**A-D**) *tg(hsp70:bmp4, cmlc2:GFP)* embryos and (E-H) wild type clutch mates at 3dpf after heat shock at timepoints between 24 and 26hpf. (**A’-H’**) lateral and (**A’’-H’’**) ventral views of the same embryos. (**A’-C’’**) Transgene embryos with induced ubiquitous expression of *bmp4* exhibit coloboma and a lateral dislocation of the lens when expression was induced between 24 and 26hpf. (**D’-H’’**) Uninduced transgene embryos and their wild type clutchmates do not develop this phenotype. (**I-L**) Whole mount in situ hybridization for *cdh6* in 30hpf embryos (prim-15). (**I**) Lateral and (**J**) dorsal view of a *tg(hsp70:bmp4, cmlc2:GFP)* embryo after heat shock at 24hpf. (**K**) Lateral and (**L**) dorsal view of a wild type embryo. *cdh6* expression in the forebrain (arrowhead) and the optic fissure (arrow) is strongly decreased by ubiquitous *bmp4* expression.

### Conclusion

In this study, we present evidence for a role of TGFβ signaling in optic fissure fusion by ECM remodeling, functionally involving *cdh6*. Our data further indicate a requirement of local inhibition of BMP signaling within the optic fissure margins during this process. This is facilitated by the TGFβ dependent induction of BMP antagonists in the margins of the optic fissure.

### Experimental Procedures

#### Zebrafish husbandry

Zebrafish (*Danio rerio*) were kept as closed stocks in accordance with local animal welfare law. The facility is under supervision of the local representative of the animal welfare agency. The fish were maintained in a constant recirculating system at 28°C on a 12h light/12h dark cycle. Fish lines used in this study were created in the AB wild type strain.

#### Gene knock down

We used three different Morpholino oligonucleotides (Gene Tools Llc) with 3’-fluorescein labels. A standard control MO (5’-CCT CTT ACC TCA GTT ACA ATT TAT A-3’), *cdh6* MO1 (5’-AAG AAG TAC AAT CCA AGT CCT CAT C-3’), and *cdh6* MO2 (5’-TCC GCT CTT AGG GTG TCT TAC AGG G-3’, both Kubota et al., 2007; Liu et al., 2008). MOs were microinjected into the cytoplasm of zebrafish zygotes at concentrations of 750μM (control MO and *cdh6* MO1) and 500μM (*cdh6* MO2). The desired injection volume was calibrated using a slide with 0.01mm scale (Bresser, Germany). The phenotype of morphants was assessed at 3dpf using a Nikon SMZ18 stereomicroscope, followed by fixation in 4%PFA for further analysis.

#### Heat shocks and controls

*tg(hsp70:bmp4, cmlc2:eGFP)* eggs were kept in a petri dish at 28°C after fertilization. To induce *bmp4* expression, 21-26hpf embryos were transferred to a 1.5ml reaction tube and incubated for 1h at 37°C in a heating block. Afterwards, they were returned to a dish at 28°C. Embryos were fixed with 4%PFA at 30hpf for in situ hybridization and at 3dpf for morphological analysis.

We used *tg(hsp70:bmp4, cmlc2:eGFP)* embryos which were not heat shocked as controls, as well as heat shocked wild type siblings from the same clutch of eggs.

#### Transgenic zebrafish

Plasmid DNA containing Smad binding elements (SBE) in combination with a minimal promoter, were kindly provided by Peter tenDijke ((CAGA)_12_ MLP Luc). Here repetitive SBEs derived from the promoter of the human plasminogen activator inhibitor gene (Dennler et al., 1998) were used to drive a luciferase gene.

We cloned the SBEs with the minimal promoter into a Gateway 5’ entry vector (Invitrogen). A multisite Gateway reaction (Kwan et al., 2007) was subsequently performed resulting in an SBE driven GFPcaax construct (SBE:GFPcaax). A Zebrafish line was generated with SB (sleeping beauty) transgenesis according to Kirchmaier et al (Kirchmaier et al., 2013). Shh reporter zebrafish were generated according to Schwend et al (Schwend et al., 2010). The plasmid was kindly provided by Sara Ahlgren.

We assembled the expression construct for *Tg(hsp70:bmp4, cmlc2:eGFP)* in a Gateway reaction, using a Tol2 destination vector including *cmlc2:eGFP* (Kwan et al., 2007), a 5’Entry vector containing the *hsp70* promotor, a pENTR D-TOPO (ThermoFisher Scientific) vector containing the CDS of *bmp4* (Heermann et al., 2015) and a 3’Entry vector with a polyadenylation site (Kwan et al., 2007). The construct (10ng/μl) was injected into wild type zebrafish zygotes together with Tol2 transposase mRNA (7ng/μl). Embryos with strong GFP expression in the heart were selected as founders. Lines were kept in closed stocks and validated in every generation.

#### Drug treatments

Zebrafish embryos were treated with SB431542 (8*μ*l/ml) to inhibit TGFβ mediated signaling. The substance was dissolved in DMSO (10 mmol stock). Controls were treated with equally concentrated DMSO without the inhibitor.

#### Microscopy

Signaling reporter fish were imaged with a Leica SP5 setup, samples mounted in glass bottom dishes (MaTek). For time-lapse imaging embryos were embedded in 1% low melting agarose covered with zebrafish medium an anesthetized with tricain. Left and right eyes were used and oriented to fit the standard views. A stereomicroscope (Olympus/ Nikon) was used for recording brightfield and fluorescent images of TGFβ signaling reporter fish. Whole mount in situ hybridizations were recorded with a stereomicroscope (Olympus) as well as an upright microscope (Zeiss). For time-lapse imaging embryos were embedded in 1% low melting agarose covered with zebrafish medium an anesthetized with tricain. Left and right eyes were used and oriented to fit the standard views.

#### Whole mount in situ hybridization

Whole mount in situ hybridizations were performed according to Quiring et al. (Quiring et al., 2004).

#### Mice

For this study TGFb2+/- (Sanford et al., 1997) and GDNF+/- (Pichel et al., 1996) mice were used for breeding. Timed matings were performed overnight and the day on which a vaginal plug was visible in the morning was considered as day 0.5. Analyses were restricted to embryonic stages because of perinatal lethality of the individual mutants. For analysis of embryonic tissue, the mother was sacrificed and the embryos were collected by caesarean sectio. All of the experiments were performed in agreement with the ethical committees. Genotyping was performed according to Rahhal, Heermann et al., 2009.

#### Histological analysis

Tissue was processed for paraffin sectioning. Frontal sections of control and TGFb2 - /- embryos were performed and stained with haematoxylin and eosin.

#### Microarray data

RNA was extracted from whole eyes of E13.5 embryos (controls and TGFb2-/-(GDNF-/-) respectively). RNA was reverse transcribed, amplified and loaded on Agilent one-color microarray chips. Experiments were performed in triplicates.

Further analysis was performed using R (R Core Team, 2014) and the bioconductor packages Agi4x44PreProcess, limma and mgug4122a.db as annotation database. For background correction we used the following parameters: BGmethod = ”half”, NORM-method = ”quantile”, foreground = ”MeanSignal”, background = ”BGMedianSignal” and offset = 50. The probes were filtered using the recommended thresholds and afterwards the replicated non-control probes were summarized. Then the method *lmFit* was used to fit a linear model on the arrays. Finally, the differential expression statistics were computed using the methods *eBayes*. Next only those genes with fold change higher than 1.5 were considered, then a multiple comparison correction was performed on the p-values using the BH (Benjamini & Hochberg) method. The genes with corrected p-value lower than 0.05 were defined as significantly differentially expressed genes.

#### Functional analysis of gene sets

We used the tool gProfiler (J. Reimand et al., 2016) (<u>http://biit.cs.ut.ee/gprofiler/)</u>) version 6.7 to find enriched terms on the set of significantly down-regulated genes from the mouse arrays. We provided the official gene symbol of these genes as input and used the default set of databases.

**Table 1:** Table: Results of *cdh6* MO injection. The table shows combined numbers of 3 independent experiments for each condition.

| condition | coloboma | ambiguous<sup>a</sup> | severe defects | total N |
| --- | --- | --- | --- | --- |
| uninjected | 0 (0%) | 1 (0.9%) | 3 (2.7%) | 113 |
| control MO | 0 (0%) | 0 (0%) | 1 (1.3%) | 76 |
| *cdh6* MO1 (11.5ng) | 30 (31.6%) | 19 (20%) | 24 (25.3%) | 95 |
| *cdh6* MO2 (4.5ng) | 40 (50.6%) | 26 (32.9%) | 1 (1.3%) | 79 |

<sup>a</sup> abnormal pigmentation of the ventral retina

## Acknowledgements

We want to thank Rolf Zeller for making Gremlin KO mice available as well as for scientific discussion. We also want to thank Gabriela Salinas-Riester and the TAL of the University Göttingen. We also thank Sara Ahlgren for generously making the shh-reporter construct available and Peter tenDijke for the luciferase construct on which our new TGFβ reporter zebrafish is based. We thank Lea Schertel and Lidia Koschny for great technical assistance and Burkhard Höckendorf for constructive input and help generating the TGFβ reporter line and Hannes Voßfeld for assistance with the mouse mutants. The project was in part funded by the Deutsche Forschungsgemeinschaft (project: Molecular mechanisms regulating optic fissure closure)

## Author contributions

MK: experiments, JM: analysis of Microarray data, PE: experiments, ER and BR: TGFβ KO mouse acquisition, AZ: sent fixed embryonic heads (Grem1 KO and controls) for analyses, KK and JW: helped conceiving the study and acquired funding, SH: conceived the study, experiments, wrote the manuscript

The authors declare that they do not have a conflict of interest.

---

**Figure S1:** Establishment of a TGFβ signaling reporter in zebrafish (**A**) construct generated for the TGFβ signaling reporter. Multimerized Smad Binding Elements (SBEs) in combination with a minimal promoter (MP) drive membrane localized GFP (GFPcaax). (**B**) Brightfield, fluorescent and merged images of a zebrafish larvae expressing the TGFβ signaling reporter construct (21.5 hpf). Note the expression domains in the forebrain (encircled) and the tail. (**C**) In order to test the functionality, de-chorionated zebrafish larvae expressing the TGFβ signaling construct were exposed to a TGFb signaling inhibitor (SB431542). The inhibitor suppressed the activity of the TGFb signaling reporter dramatically (recorded at postembryonic stages). Please see the DMSO treated fish as control.

**Figure S2:** Early ubiquitous *bmp4* expression causes defects in optic cup morphogenesis. (A-F) Gross morphology of (A-C) *tg(hsp70:bmp4, cmlc2:GFP)* embryos and (D-F) wild type clutch mates at 3.5dpf after heat shock at timepoints between 21 and 23hpf. (A’-F’) lateral and (A’’-F’’) ventral views of the same embryos. (A’-C’’) Transgene embryos with ubiquitous expression of *bmp4* induced between 21 and 23hpf show coloboma caused by impaired optic cup morphogenesis as revealed by the lack of ventral retina and increased size of the temporal retinal domain. (D’-F’’) Uninduced transgene embryos and their wild type clutchmates do not develop this phenotype.

